# Inter-brain similarity and connectivity based on EEG hyperscanning during patient‒acupuncturist interactions

**DOI:** 10.1101/2025.04.29.651215

**Authors:** Shuyong Jia, Qi Liu, Na Tu, Yuning Qin, Shenjun Wang, Qiuyue Lyu, Yuhan Liu, Zixin Huo, Xiaojing Song, Shuyou Wang, Weibo Zhang, Liyun He, Yi Guo, Guangjun Wang

**Affiliations:** Institute of Acupuncture and Moxibustion, China Academy of Chinese Medical Sciences; Beijing 100700, China; Tianjin University of Traditional Chinese Medicine; Tianjin 301617, China; Institute of Basic Research in Clinical Medicine, China Academy of Chinese Medical Sciences; Beijing 100700, China; National Clinical Research Center for Chinese Medicine Acupuncture and Moxibustion; Tianjin 300381, China

**Keywords:** Electroencephalogram (EEG) hyperscanning, Inter-brain similarity, Inter-brain connectivity, Patient–acupuncturist interactions, High-frequency oscillations (HFOs)

## Abstract

The study of patient‒clinician relationships is a vital branch of interpersonal interaction research. Acupuncture attaches great importance to positive patient‒clinician interactions; thus, it serves as a typical example of patient‒clinician interaction research. Based on electroencephalography (EEG) hyperscanning, this study explored inter-brain similarity based on spatial correlations between two brains as well as inter-brain connectivity of different regions through different frequency bands. Regarding inter-brain similarity, both acupuncture and sham acupuncture significantly decreased spatial correlation during certain sessions during acupuncture. Regarding inter-brain connectivity, acupuncture enhanced not only the connectivity of the beta frequency band in the right temporoparietal regions but also the connectivity of the fast-ripple frequency band in the right and left temporoparietal regions. These results suggest that inter-brain similarity should be emphasized in patient–clinician relationship research. Moreover, high-frequency oscillations (HFOs) potentially play an important role in inter-brain connectivity in addition to the classical EEG low-frequency bands.

## 1 INTRODUCTION

Acupuncture has been widely applied for more than three thousand years, is one of the most popular forms of alternative and complementary medicine in the world, and is widely used to alleviate or cure various diseases^1,2^. There is a strong placebo effect that is an essential component of acupuncture efficacy during the process of traditional Chinese acupuncture therapy^3,4^. Many ancient studies and modern studies have confirmed that full and active participation and a high degree of cooperation between the acupuncturist and patient enhance the placebo effect, leading to a better therapeutic effect^5–7^. Therefore, patient‒acupuncturist relationships represent an important part of all acupuncture therapeutic environments, and a positive relationship during patient‒ acupuncturist interactions plays a significant role in the effectiveness of treatment outcomes.

Patient–acupuncturist relationships during interactions involve social psychology, which is the study of how people’s thoughts, feelings, and behaviours are influenced by the real or imagined presence of other people^8,9^. Hyperscanning is a common and effective method used to study social psychology. Researchers can use hyperscanning to track dynamic social interactions, in which multiple people engage in different forms of social interaction in real time^10^. For example, recent functional magnetic resonance imaging (fMRI) and electroencephalography (EEG) studies of patients and clinicians during interactions (i.e., hyperscanning) have explored the brain processes underpinning the social modulation of pain. These studies revealed that both behavioural and brain responses across the patient‒clinician dyad were significantly affected by the interaction style^11^, both empathy and supportive care can reduce pain intensity^12^, patient‒clinician concordance in brain activity represents a potentially key mechanism mediating patient analgesia^13^ and patient‒clinician concordance of the insula plays an important role in mediating the directional dynamics of pain-directed facial communication during therapeutic encounters^14^.

Among hyperscanning studies of acupuncture, Ellingsen et al. performed fMRI hyperscanning of 37 patient‒acupuncturist dyads who interacted via live video with each other while acupuncturists treated evoked pain in patients with chronic pain^13^. These researchers reported that dynamic patient‒acupuncturist concordance in brain activity implicated in social mirroring and theory of mind was increased after the clinical interaction. In this study, acupuncture was practised in an environment far from a real clinical environment. Therefore, Chen et al. used functional near-infrared spectroscopy (fNIRS) hyperscanning to explore the inter-brain mechanism of patient‒ acupuncturist dyads in a more realistic clinical setting^15^. By simultaneously recording the neural responses of simulated patients and acupuncturists during acupuncture stimulation in each group and analysing the results of the acupuncture and sham acupuncture groups, Chen et al. reported an increase in inter-brain neural synchronization (INS) in the prefrontal cortex (PFC) between the simulated patient and the acupuncturist during acupuncture but not sham acupuncture stimuli. This study provides a good example for investigating the neurobiological mechanism of patient‒ acupuncturist interactions during acupuncture stimuli in a natural clinical setting.

However, the temporal resolution of fNIRS is not sufficient. Patient–acupuncturist relationships involve a wide range of psychological activities that incorporate different cognitive and brain processes. Certain elementary mental processes, such as cognitive processing, can often occur quite rapidly, requiring the use of measures with very high temporal resolution to study them^16,17^. In addition, fNIRS only explores the functional activation of the human cerebral cortex by measuring brain tissue concentration changes in oxygenated and deoxygenated haemoglobin following neuronal activation and does not provide a direct measurement that reflects brain neural activity^18,19^.

Nearly all neuroimaging techniques, such as fMRI, EEG and fNIRS, can be used for hyperscanning^20^. However, as a powerful and popular technique of neurophysiology, EEG is increasingly used in acupuncture research and seems to be more suitable for exploring patient‒acupuncturist relationships during acupuncture interactions^21,22^. First, EEG is inexpensive and noninvasive. In addition, compared with fNIRS, EEG has a greater temporal resolution, which can provide a more fine-tuned account of neural content and timing during patient‒acupuncturist interactions and more quickly capture the momentary changes in the acupuncture signal in the brain; moreover, it directly reflects the neurophysiological activity of both the patient and the acupuncturist by measuring neuroelectric signals in the brain. Anzolin et al. performed EEG hyperscanning experiments to explore the brain and behavioural mechanisms of the patient‒acupuncturist relationship while engaging in acupuncture treatment^11^. However, because only the EEG signals of the patient and acupuncturist during the electroacupuncture treatment phase were analysed separately, this study differs from a real acupuncture treatment situation with the needle insertion, needle manipulation and needle withdrawal sessions^23^ and does not take full advantage of the synchronization of EEG hyperscanning to computationally study the brain similarity, connectivity and synchronization between the patient and acupuncturist.

Different sessions during acupuncture, such as needle insertion, needle manipulation and needle withdrawal, involve different patient‒acupuncturist interactions. The current study aimed to establish patient–acupuncturist EEG hyperscanning with simultaneous recordings of patient–acupuncturist interactions during different sessions in acupuncture and investigate the differences in inter-brain similarity and connectivity in the patient–acupuncturist dyad between acupuncture and sham acupuncture and the changes in that during different patient–acupuncturist interaction sessions. We hypothesized that acupuncture would change the inter-brain similarity and connectivity during patient‒acupuncturist interactions and that the change in inter-brain similarity and connectivity would occur during both acupuncture and sham acupuncture. In addition, some studies have suggested that high-frequency oscillations (HFOs) are involved in certain brain activities^24^; however, how HFOs function in interpersonal interactions is not well understood. Therefore, based on the present study of patient– acupuncturist inter-brain similarity and connectivity in low-frequency bands, we also aimed to further explore whether HFOs are involved in patient‒acupuncturist interactions. We hypothesized that acupuncture and sham acupuncture affect inter-brain similarity and connectivity in HFOs. These results enhance our understanding of patient‒acupuncturist interactions and provide a new approach for exploring the central mechanism of the effects of acupuncture and sham acupuncture.

## 2 MATERIALS AND METHODS

### 2.1 Participants

Sixty healthy subjects (simulated patients) were recruited for this study from August 2023 to November 2023. All participants were right-handed and had no history of neurological or psychiatric disorders. Additionally, participants had normal or corrected-to-normal vision. These simulated patients were randomly assigned to either the acupuncture group or the sham acupuncture group at a 1:1 ratio. One healthy licenced male acupuncturist was recruited. This acupuncturist performed acupuncture on all 60 subjects, resulting in 60 patient–acupuncturist dyads in total.

All the participants signed an informed consent form before the experiment. All the experimental procedures were conducted in accordance with the Declaration of Helsinki^25^. Ethical approval was granted by the Ethics Committee of the Institute of Acupuncture and Moxibustion, China Academy of Chinese Medical Sciences.

### 2.2 Experimental procedures

Throughout the experiment, all dyads sat during the entire acupuncture session in a quiet room. The experimental setup is shown in Fig. 1a, b. A separator was placed in between the dyad to block their ability to communicate or make eye contact with each other, but the right arm of the simulated patient was able to pass through the separator to receive the acupuncture. Simulated patients seated 80 cm from a computer screen were presented with a red fixation cross and instructions with E-Prime 2.0. The acupuncturist was presented with a red fixation cross at the centre of a white background placed 70 cm in front of him. In addition, the task and the whole experimental procedure included the following four main sessions (Fig. 1e).

1. The baseline: First, the computer screen provided instructions about the experimental requirements to the patient. The patient had to press the numeric keypad with the left hand to indicate that he or she fully understood the instructions (marked as “11”). Next, voice instructions were provided through a speaker to tell the acupuncturist that the patient was ready (marked as “21”). Then, a red fixation cross appeared at the centre of the screen for 6 min. The patient stared at the cross during this period, and the acupuncturist stared at the same cross in front of him.
2. Needle insertion: Second, the computer screen provided instructions to tell the simulated patient to prepare for needle insertion. After 10 s, voice instructions altered the acupuncturist to start inserting the needle (marked as “22”). When the acupuncturist inserted the needle, the patient was instructed to press the numeric keypad with the left hand to indicate the feeling. If there was a sensation of needle insertion, the keypad was pressed (marked as “12”). Then, the screen displayed the next interface. If there was no sensation of needle insertion, the keypad was not pressed. Then, the screen displayed the next interface 30 s after the “22” mark. The next interface displayed a red fixation cross for 6 min.
3. Needle manipulation: Third, instructions appeared on the computer screen to tell the simulated patient to prepare for the needle manipulation. After 10 s, voice instructions were provided to the acupuncturist to start manipulating the needle (marked as “23”). When the acupuncturist manipulated the needle, the patient was instructed to press the numeric keypad with the left hand according to his or her feelings. If the patient felt *deqi* sensation, the patient pressed the keypad (marked as “13”). Then, the acupuncturist was verbally given instructions to stop the needle manipulation (marked as “24”). Then, the screen displayed the next interface. If there was no *deqi* sensation, the keypad was not pressed. Then, 30 s after the “23” mark, the acupuncturist was given voice instructions to stop needle manipulation (marked as “24”). Then, the next interface appeared. The next interface displayed a red fixation cross for 10 min.
4. Needle withdrawal: Finally, the computer screen provided instructions to tell the simulated patient to prepare for needle withdrawal. After 10 s, the acupuncturist was verbally instructed to withdraw the needle (marked as “25”). When the acupuncturist withdrew the needle, the patient had to press the keypad with the left hand based on the feelings. If there was a sensation of needle withdrawal, the keypad was pressed (marked as “14”). Then, the screen displayed the next interface. If there was no sensation of needle withdrawal, the keypad was not pressed. Then, 15 s after the “25” mark, the screen displayed the next interface. The next interface displayed a red fixation cross for 6 min.

**Fig. 1.**
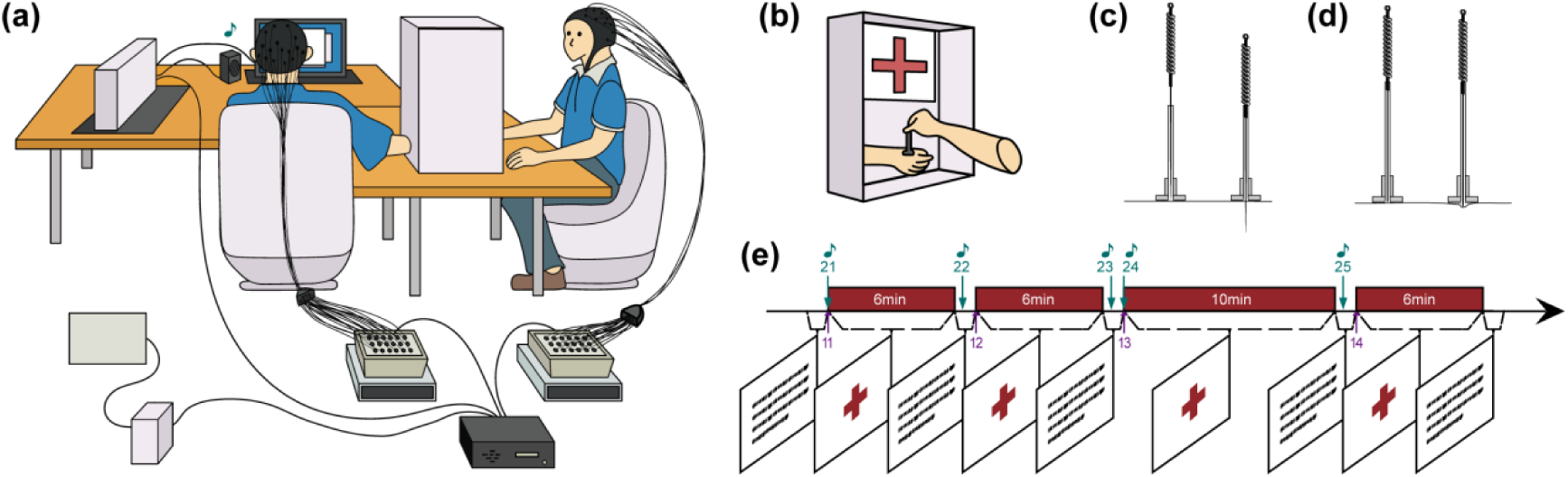
Experimental design and task procedures. (a) and (b) Experimental setup. (c) The mechanism of a real needle. After the real needle passes through the tube and touches the skin (left), it penetrates the skin with a maximum depth of 13.5 mm (right). (d) The mechanism of a sham needle. After the sham needle passes through the tube and touches the skin (left), it hits the skin surface at a maximum depth of 1.5 mm without penetration (right). (e) Acupuncture experimental paradigm. Four main EEG recordings were obtained during the experiment: baseline (6 min), post-needle insertion (6 min), post-needle manipulation (10 min) and post-needle withdrawal (6 min). Different green numbers and arrows indicate different behaviours of the acupuncturist. Different purple numbers and arrows indicate different behaviours of the simulated patient. The green notes indicate voice instructions from a speaker. Parallelograms indicate instructions from a computer screen.

### 2.3 Acupuncture methods

Details of the acupuncture method are shown in Fig. 1c, d. The devices used for both acupuncture and sham acupuncture are composed of two parts: a needle and a plastic tube connected with a sticky opaque base on the skin. The needles used in this study were 0.30 mm × 50 mm in size. In addition, the diameter of the needle handle is larger than that of the plastic tube. The tip of the real needle was sharp. For the sham needle, the tip was blunt.

All participants received manual acupuncture at the LI 4 (Hegu) acupuncture point situated between the first and second metacarpal bones on the dorsal side of the right hand^26^. After skin disinfection, the acupuncture needles were inserted into the deepest part. Small, equal needle manipulations of lifting and thrusting were subsequently performed on all needles for 30 s after 6 min of needle insertion. If the simulated patients experienced *deqi* sensations, the needle manipulations were stopped immediately. Next, all needles were withdrawn after 10 min.

In addition, prior to acupuncture, all the simulated patients received video training, which helped them assess whether *deqi* sensations were experienced during needle manipulation. After acupuncture, the patients were asked to complete the visual analogue scale (VAS) behaviour questionnaire to evaluate *deqi* sensations^22,27,28^.

### 2.4 EEG data acquisition

The EEG data were recorded with two 32-electrode EEG recording caps (Easycap GmbH, Germany) and two 24-bit EEG amplifiers (NeurOne EXG, Bittium, Finland) in DC mode at a sampling rate of 20 kHz^29^. The two amplifiers were connected to the same main unit via two optical cables. The recording electrodes were located at standard locations following the international 10-20 system^30^. FCz and AFz served as the reference and ground electrodes, respectively^31^. The 32 active electrodes used for the experiment are shown in Fig. 4a. The EEG signal was digitized via NeurOne software. The scalps of the subjects were prepared for EEG recording using a conductive gel (Quick-Gel, Neuromedical Supplies®)^32^. The electrode impedance was generally reduced to below 5 kΩ prior to data collection. The subjects were instructed to relax and avoid any specific mental operations.

Moreover, the NeurOne system was used simultaneously to record a pair of vertical electrooculograms (VEOGs) and a pair of horizontal electrooculograms (HEOGs) at supra/suborbital and external canthi sites, respectively, as these data may be useful for removing ocular artefacts during a later processing step^33^. Finally, two cutaneous electrodes were connected to the same system to acquire the ECG data.

### 2.5 EEG data analysis

#### 2.5.1 Preprocessing

The EEG data were preprocessed via the EEGLAB toolbox^34^ in MATLAB (version 2022b; MathWorks Inc.). With respect to the EEG data at low-frequency bands, the recorded data were first high-pass filtered at 1 Hz and low-pass filtered at 70 Hz, after which they were resampled at 500 Hz. The data were subsequently separated into two recordings corresponding to each participant. The function *cleanLineNoise*^35^ was subsequently used to remove line noise. Next, reference electrode standardization technique (REST)^36^ was applied to reset the place of reference electrode. Then, bad channels were interpolated. Next, an artifact subspace reconstruction (ASR) algorithm^37–39^ adapted for EEGLAB software (clean_rawdata plugin) was implemented to remove high-amplitude artifacts (for example, those stemming from eye blinks, muscle, and sensor motion) from the EEG signal. The independent component analysis (ICA) technique was subsequently applied to the data obtained from each participant. Next, an equivalent dipole current source was fit to each IC via the DIPFIT toolbox of EEGLAB^40^. Individual components accounting for blinks and saccades, heartbeats, muscle artifacts or line noise were subsequently removed from the data based on the exclusion criteria outlined by Perez et al.^39^. Finally, the EEG data of 300 s, 300 s, 540 s, and 300 s were extracted and saved from 30 s after the voice-instructed markers “21”, “22”, “24”, and “25”, respectively, indicating the EEG data at baseline, after needle insertion, after needle manipulation, and after needle withdrawal for each subject.

For the EEG data at HFOs, the recorded data were first resampled at 2000 Hz. Next, the data were separated into two recordings corresponding to each participant and further divided into four epochs of EEG data in the same manner as described for low-frequency bands.

#### 2.5.2 Spatial correlation analysis

Spatial correlation is an amplitude-based connectivity measure used to estimate the connectivity between brains at the global level. The formula used to calculate the spatial correlation is as follows.

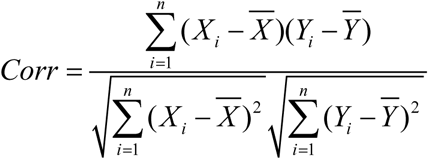

where *Corr* represents the spatial correlation at each sample point. *X_i_* is the amplitude for the *i*’th channel of the simulated patient at each sample point. 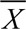 is the average amplitude for all channels of the simulated patient at each sample point. *Y_i_* is the amplitude for the *i*’th channel of the acupuncturist at each sample point. 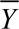 is the average amplitude for all channels of the acupuncturist at each sample point. *n* is the number of channels of the signal. We used MATLAB to obtain the spatial correlations for all dyads. The preprocessed clean data were imported first. Next, for the low-frequency bands in the EEG data, the data were filtered into four frequency bands: theta (4–8 Hz), alpha (8–12 Hz), beta (12–30 Hz) and 4–30 Hz low-frequency band^41^. For EEG data at HFOs, the data were filtered into four frequency bands: high-gamma (80– 150 Hz), ripple (150–250 Hz), fast-ripple (250–500 Hz) and 80–500 Hz high-frequency band^42^. In addition, spatial correlations between “patient” and acupuncturist brains at all sample points were calculated for each frequency band separately. The spatial correlations at all sample points were smoothed. Then, the positive spatial correlations were selected for averaging to obtain the average spatial correlation for the dyad under each frequency band. Finally, Fisher’s Z transformation was then used to normalize the spatial correlation to confirm its approximately normal distribution such that the subsequent statistical analysis could be performed logically^43^.

#### 2.5.3 Inter-brain connectivity analysis

Artifact-free EEG data acquired simultaneously from each dyad were subjected to inter-brain connectivity analysis, which provides information on synchronous activity between scalp channels and areas within each dyad. We used the Python open source software package “HyPyP”^44^ to analyse the changes in inter-brain connectivity. Measuring the phase difference between EEG channels is a method used to estimate the connectivity between channels, and the phase locking value (PLV) is one of the most popular phase-based connectivity measures^45^. However, the PLV could eventually provide spurious connections, which could be overcome using CCorr to estimate phase synchrony^45,46^. Therefore, we used CCorr in this study to estimate the phase synchrony between channels in the “patient” and acupuncturist brains.

First, the preprocessed clean data were imported. For each dyad, the EEG data from the simulated patient and acupuncturist were aligned and merged into a single file, and a total of 4 EEG data were formed for each pair. Second, the EEG data were segmented into consecutive epochs of 2 s in preparation for the hyper-connectivity analysis. Next, for EEG data at low-frequency bands, the data were filtered into three frequency bands: theta (4–8 Hz), alpha (8–12 Hz) and beta (12–30 Hz)^41^. For EEG data at HFOs, the data were filtered into three frequency bands: high gamma (80–150 Hz), ripple (150–250 Hz) and fast-ripple (250–500 Hz)^42^. The instantaneous phase was subsequently estimated using the Hilbert transform. In addition, for all possible channel pairings, CCorr values were calculated for each 2 s epoch and frequency band separately. Then, the CCorr values for all epochs within each EEG data under the same frequency band were averaged to obtain the CCorr value for each EEG data under each frequency band.

Finally, for the ROIs, i.e., frontal (Fp2, Fz, Fp1, F3, F4, F7, F8), frontal-central (Cz, FC1, FC2, C3, C4), parietal (Cp1, P3, Pz, Cp2, P4), left temporoparietal (FC5, FT9, T7, CP5, P7) and right temporoparietal (P8, CP6, T8, FT10, FC6) ^47^, the CCorr values of all possible channel pairings under the same frequency band in the same region were averaged via MATLAB to obtain the average CCorr value for the EEG data of each brain region under each frequency band. Fisher’s Z transformation was then used to normalize the CCorr values to obtain an approximately normal distribution.

#### 2.5.4 Statistical analyses

Statistical analyses were performed using IBM SPSS Statistics (version 21) and MATLAB (version 2022b). First, statistical analyses were performed to analyse the subject demographics. For the statistical comparison of categorical variables, we used the Chi-square test with Fisher’s exact test. In the case of continuous variables, Shapiro‒Wilk tests were first applied to check the normality of the variables. For Gaussian variables, an independent-sample *t*-test was applied. For non-Gaussian variables, a nonparametric test (Mann‒Whitney *U* test) was performed between the two groups.

For the percent change from baseline in the spatial correlation, given its normality, a mixed two-way ANOVA with one within-subjects factor (acupuncture session: baseline, post-needle insertion, post-needle manipulation, and post-needle withdrawal) and one between-subjects factor (group: acupuncture, sham acupuncture) was performed. The sphericity of the data was assessed using a Mauchly test. Post-hoc tests were performed with two-tailed paired or unpaired *t*-tests, depending on the factor, and false discovery rate (FDR) corrected.

In addition, for inter-brain connectivity, an independent two-sample t-test with FDR correction was used to separately assess the between-group differences in the inter-brain connectivity of channels at different sessions during acupuncture. To mitigate the effects of factors such as volume conduction, the ROIs were used to group the EEG data for further analysis. Given the normality of CCorr, a mixed two-way ANOVA was performed with the same details as the statistical analysis of spatial correlation.

## 3 RESULTS

### 3.1 Participants

We scanned a total of 60 dyads and obtained complete EEG data from 57 dyads (29 in the acupuncture group and 28 in the sham acupuncture group, with 2 incomplete dyads because of record interruption and 1 incomplete dyad because of no voice instructions). Thus, 57 patient–acupuncturist dyads were included in the final analyses. The group demographic characteristics are presented in Table 1. Participants receiving acupuncture and sham acupuncture did not differ in terms of sex (Fisher’s exact test, *p* = 0.297), height (Mann‒Whitney *U* test, *p* = 0.154), weight (Mann‒Whitney *U* test, *p* = 0.159) or age (Mann‒Whitney *U* test, *p* = 0.879). For each dyad, participants were asked to work together on acupuncture, and EEG was used to collect brain activity from both the simulated patient and the acupuncturist simultaneously (Fig. 1). During the experiment, a baseline session was followed by an acupuncture task that included three sessions: needle insertion, needle manipulation and needle withdrawal (see the Materials and methods section for details).

**Table 1.** Demographic characteristics.

| **Characteristics** | **Acupuncture**<br>**(*n*=29)** | **Sham acupuncture**<br>**(*n*=28)** | ***p* value** |
| --- | --- | --- | --- |
| Sex |  |  | 0.325 |
| Male | 7 (24.1) | 3 (10.7) |  |
| Female | 22 (75.9) | 25 (89.3) |  |
| Height (cm) | 163.0 (158.0-172.0) | 160.0 (157.3-165.0) | 0.154 |
| Weight (kg) | 60.0 (53.5-69.0) | 54.0 (50.0-60.0) | 0.159 |
| Age (years) | 24.0 (24.0-30.0) | 26.0 (23.0-27.8) | 0.879 |

Continuous variables reported as median (interquartile range). Categorical variables reported as *n* (%).

### 3.2 Spatial correlations in low-frequency bands

The spatial correlation between the patient and acupuncturist was obtained by calculating their EEG data (Fig. 2a, b) (see the Materials and methods section for details). The 4–30 Hz low-frequency band contains three frequency bands (theta, alpha and beta). Regarding the percent change from baseline in the spatial correlation at 4– 30 Hz, theta and alpha frequency bands, a significant main effect of the acupuncture session was found (mixed-design ANOVA, 4–30 Hz: F (2.77, 152.21) = 63.16, *p* < 0.001; theta: F (2.89, 158.89) = 47.02, *p* < 0.001; and alpha: F (3, 165) = 63.51, *p* < 0.001) (Fig. 2c–e). Post-hoc tests revealed a significant increase in baseline, post-needle insertion and post-needle withdrawal compared with post-needle manipulation in these three frequency bands (two-tailed paired *t*-test, baseline vs. post-needle manipulation: *p* < 0.001; post-needle insertion vs. post-needle manipulation: *p* < 0.001; post-needle manipulation vs. post-needle withdrawal: *p* < 0.001).

**Fig. 2.**
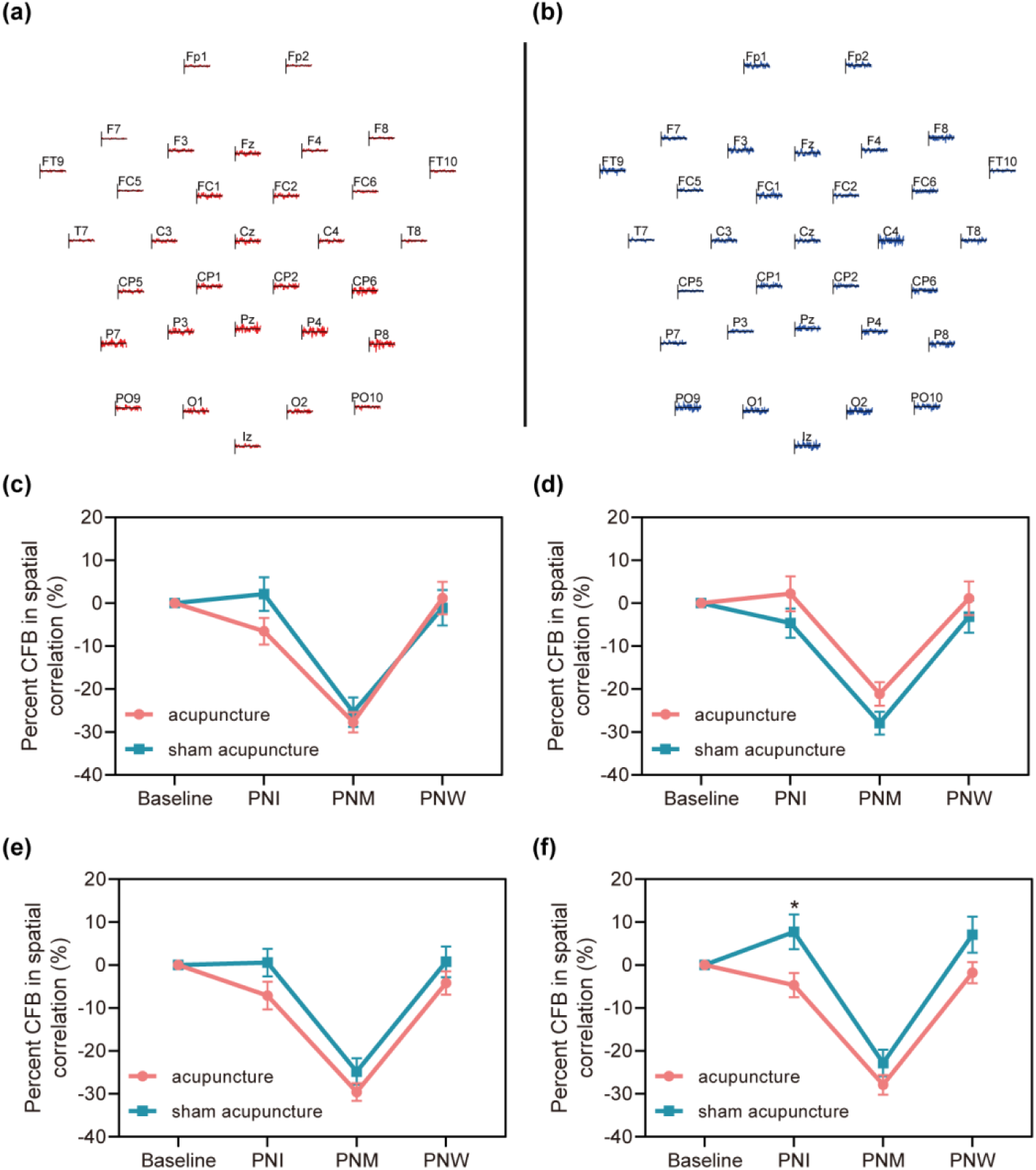
Acquisition of spatial correlation and change from baseline in spatial correlation at different sessions in low-frequency bands. (a) Example of EEG data from different channels of the simulated patient. (b) Example of EEG data from different channels of the acupuncturist. (c) Percent CFB in spatial correlation at baseline, PNI, PNM and PNW sessions in the 4–30 Hz low-frequency band. (d) Percent CFB in spatial correlation at baseline, PNI, PNM and PNW sessions in the theta frequency band (4–8 Hz). (e) Percent CFB in spatial correlation at baseline, PNI, PNM and PNW sessions in the alpha frequency band (8–12 Hz). (f) Percent CFB in spatial correlation at baseline, PNI, PNM and PNW sessions in the beta frequency band (12–30 Hz). The star indicates significant difference between the acupuncture and sham acupuncture groups. The data in (c)-(f) are plotted as the mean±SEM. Acupuncture, *n*=29; sham acupuncture, *n*=28. CFB, change from baseline; PNI, post-needle insertion; PNM, post-needle manipulation; PNW, post-needle withdrawal; SEM, standard error of the mean.

Regarding the percent change from baseline in the spatial correlation in the beta frequency band, a significant effect was found for the SESSION × GROUP interaction (mixed-design ANOVA, F (3, 165) = 2.89, *p* = 0.037) (Fig. 2f). Post-hoc tests revealed a significant decrease in the acupuncture group compared with the sham acupuncture group post-needle insertion (two-tailed unpaired *t*-test, *p* = 0.014). Additionally, similar to the other low-frequency bands, significant reductions were observed post-needle manipulation in both the acupuncture and sham acupuncture groups (two-tailed paired *t*-test, baseline vs. post-needle manipulation: *p* < 0.001; post-needle insertion vs. post-needle manipulation: *p* < 0.001; post-needle manipulation vs. post-needle withdrawal: *p* < 0.001), and compared with the baseline, a significant increase was found for post-needle insertion and post-needle withdrawal in the sham acupuncture group (two-tailed paired *t*-test, baseline vs. post-needle insertion: *p* = 0.032 and baseline vs. post-needle withdrawal: *p* = 0.045, respectively). These results revealed a decrease in spatial correlation in low-frequency bands after needle manipulation and a change in spatial correlation in the beta frequency band.

### 3.3 Circular correlation coefficients in low-frequency bands

Although the spatial correlations described above revealed a decrease in inter-brain similarity between the simulated patient and acupuncturist in the beta frequency band after needle insertion at the global level, the circular correlation coefficient (CCorr) is an alternative analytic strategy is used analyse the EEG signals at the channel level.

The CCorr values for all EEG channel pairings were first separately extracted in three low-frequency bands (theta, alpha and beta). These values could be used to plot connectivity matrices and assess the differences between the acupuncture and sham acupuncture groups after needle insertion, manipulation and withdrawal. Matrices and results were shown in supplementary results and Figures S1-S6. To mitigate the effects of factors such as volume conduction on the above results, regions of interest (ROIs) were used to group the EEG data, and further analyses were performed (Fig. 3a). Figure 3 and Figures S7‒S10 report the percent change from baseline in CCorr values for the two groups (post-needle insertion, post-needle manipulation, and post-needle withdrawal).

**Fig. 3.**
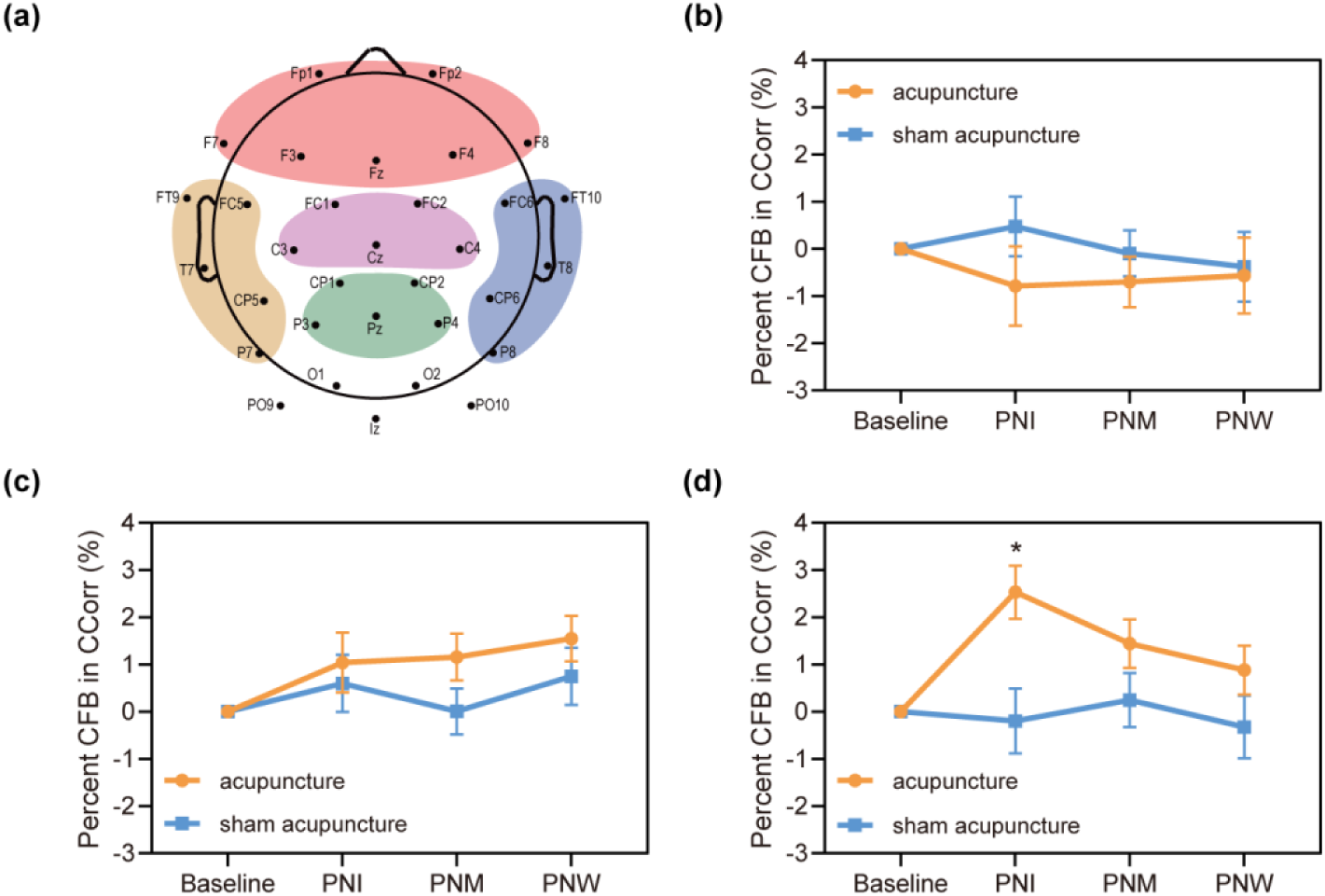
ROIs and change from baseline in CCorr in the right temporoparietal regions at different sessions in low-frequency bands. (a) The ROIs. The regions marked in the red area represent the frontal regions, the purple area represent the frontal-central regions, the green area represent the parietal regions, the brown area represent the left temporoparietal regions, and the blue area represent the right temporoparietal regions. (b) Percent CFB in CCorr in the theta frequency band (4–8 Hz) in the right temporoparietal regions at baseline, PNI, PNM and PNW sessions. (c) Percent CFB in CCorr in the alpha frequency band (8–12 Hz) in the right temporoparietal regions at baseline, PNI, PNM and PNW sessions. (d) Percent CFB in CCorr in the beta frequency band (12–30 Hz) in the right temporoparietal regions at baseline, PNI, PNM and PNW sessions. The star indicates significant difference between the acupuncture and sham acupuncture groups. The data in (b)-(d) are plotted as the mean±SEM. Acupuncture, *n*=29; sham acupuncture, *n*=28. CFB, change from baseline; PNI, post-needle insertion; PNM, post-needle manipulation; PNW, post-needle withdrawal; SEM, standard error of the mean.

The right temporoparietal region plays an important role in patient‒acupuncturist interactions during acupuncture. ANOVA results revealed a significant effect of the SESSION × GROUP interaction in the right temporoparietal regions in the beta frequency band (mixed-design ANOVA, F (3, 165) = 3.61, *p* = 0.015) (Fig. 3d). Post-hoc tests revealed a significant increase in the acupuncture group compared with the sham acupuncture group at the post-needle insertion session (two-tailed unpaired *t*-test, *p* = 0.003). Post-hoc tests also revealed a significant increase in the acupuncture group at post-needle insertion (two-tailed paired *t*-test, baseline vs. post-needle insertion: *p* < 0.001 and post-needle insertion vs. post-needle withdrawal: *p* = 0.009, respectively) and post-needle manipulation (two-tailed paired *t*-test, baseline vs. post-needle manipulation, *p* = 0.01) but not in the sham group (two-tailed paired *t*-test, *p* > 0.05). In addition, a significant main effect of the acupuncture session in the alpha frequency band was found in the right temporoparietal regions (mixed-design ANOVA, F (3,165) = 2.72, *p* = 0.046), and post-hoc tests revealed a significant increase in the alpha frequency band post-needle withdrawal compared with baseline (Fig. 3c). No significant differences or changes in the theta frequency band were found in the right temporoparietal regions (Fig. 3b).

Moreover, a significant effect was found for the SESSION × GROUP interaction on the beta frequency band in the parietal regions (mixed-design ANOVA, F (2.8, 154.17) = 4.12, *p* = 0.009) (Fig. S7d). Post-hoc tests revealed a significant increase in the acupuncture group compared with the sham acupuncture group at post-needle withdrawal (two-tailed unpaired *t*-test, *p* = 0.002). Post-hoc tests also revealed a significant increase in the acupuncture group at post-needle withdrawal compared with baseline, post-needle insertion and post-needle manipulation (two-tailed paired *t*-test, baseline vs. post-needle withdrawal: *p* = 0.002; post-needle insertion vs. post-needle withdrawal: *p* = 0.003; post-needle manipulation vs. post-needle withdrawal: *p* = 0.023) and a significant increase in the sham acupuncture group at post-needle manipulation compared with post-needle insertion and post-needle withdrawal (two-tailed paired *t*-test, post-needle insertion vs. post-needle manipulation: *p* = 0.016 and post-needle manipulation vs. post-needle withdrawal: *p* = 0.021, respectively). No significant difference was found in other frequency bands in the parietal regions (Fig. S7b, c). Similarly, no significant differences or changes in the percent change from baseline in CCorr values in the three frequency bands at the other ROIs were found (Figs. S8‒S10).

### 3.4 Spatial correlations in HFOs

To further explore the effects of acupuncture on inter-brain similarity and connectivity in high-frequency bands between the simulated patients and acupuncturists, we performed EEG to assess the high-frequency bands using mixed-design ANOVA statistical tests for spatial correlations.

The 80–500 Hz high-frequency band contains three bands (high-gamma, ripple and fast-ripple). Regarding the percent change from baseline in the spatial correlation at 80–500 Hz, significant main effects of the acupuncture session on high-gamma, ripple and fast-ripple frequency bands was found (mixed-design ANOVA, 80–500 Hz: F (2.16, 118.7) = 24.66, *p* < 0.001; high-gamma: F (2.74, 150.75) = 61.19, *p* < 0.001; ripple: F (2.53, 139.03) = 39.63, *p* < 0.001; fast-ripple: F (2.18, 120.14) = 8.01, *p* < 0.001) (Fig. 4a‒d). Similar to the low-frequency bands, post-hoc tests revealed a significant increase in all high-frequency bands at baseline, post-needle insertion and post-needle withdrawal compared with post-needle manipulation (two-tailed paired *t*-test, baseline vs. post-needle manipulation: *p* < 0.01; post-needle insertion vs. post-needle manipulation: *p* < 0.001; post-needle manipulation vs. post-needle withdrawal: *p* < 0.001).

**Fig. 4.**
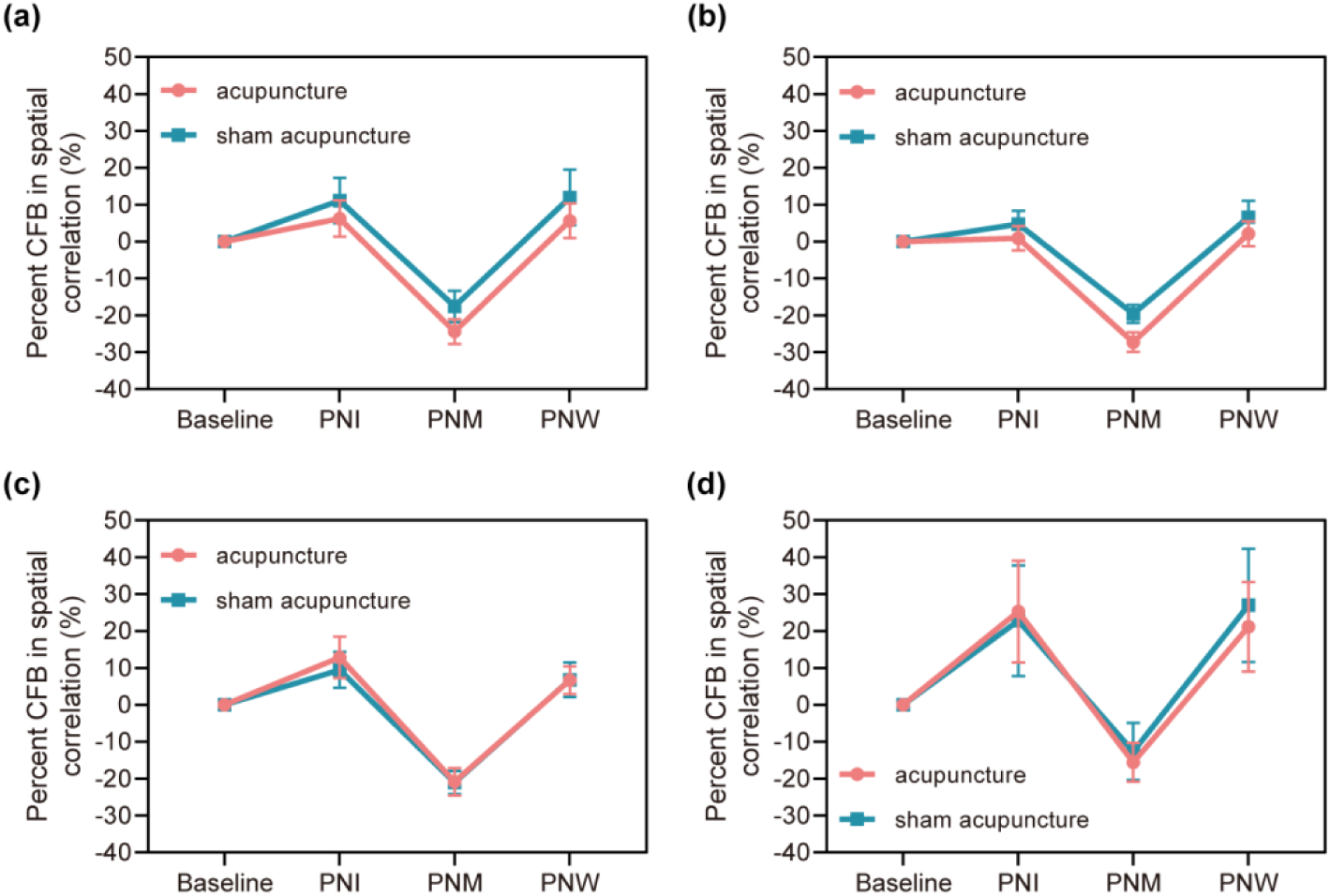
Acquisition of spatial correlation and change from baseline in spatial correlation at different sessions in high-frequency bands. (a) Percent CFB in spatial correlation at baseline, PNI, PNM and PNW sessions in the 80–500 Hz high-frequency band. (b) Percent CFB in spatial correlation at baseline, PNI, PNM and PNW sessions in the high-gamma frequency band (80–150 Hz). (c) Percent CFB in spatial correlation at baseline, PNI, PNM and PNW sessions in the ripple frequency band (150-250 Hz). (d) Percent CFB in spatial correlation at baseline, PNI, PNM and PNW sessions in the fast-ripple frequency band (250-500 Hz). The data in (a)-(d) are plotted as the mean±SEM. Acupuncture, *n*=29; sham acupuncture, *n*=28. CFB, change from baseline; PNI, post-needle insertion; PNM, post-needle manipulation; PNW, post-needle withdrawal; SEM, standard error of the mean.

Additionally, post-hoc tests revealed a significant increase in the 80–500 Hz high-frequency band at post-needle insertion compared with baseline (two-tailed paired *t*-test, *p* = 0.03) and a significant increase in the ripple frequency band at post-needle insertion and post-needle withdrawal compared with baseline (two-tailed paired *t*-test, baseline vs. post-needle insertion: *p* = 0.004 and baseline vs. post-needle withdrawal: *p* = 0.029, respectively) and the fast-ripple frequency band (two-tailed paired *t*-test, baseline vs. post-needle insertion: *p* = 0.022 and baseline vs. post-needle withdrawal: *p* = 0.017, respectively). These results reveal a decrease in spatial correlation in high-frequency bands after needle manipulation and a change in spatial correlation in high-frequency bands.

### 3.5 Circular correlation coefficients in HFOs

Although the spatial correlations described above revealed a change in the inter-brain similarity in high-frequency bands between the simulated patients and acupuncturists during the acupuncture sessions but no difference between the acupuncture and sham acupuncture groups, the CCorr was used to further assess the differences between the two groups.

The CCorr values for all EEG channel pairings were first separately extracted in three high-frequency bands (high-gamma, ripple and fast-ripple). These values could be used to plot connectivity matrices and assess the differences between the acupuncture and sham acupuncture groups after needle insertion, manipulation and withdrawal. Matrices and results were shown in supplementary results and Figures S11-S16. To mitigate the effects of factors such as volume conduction on the above results, the ROIs were used to group the EEG data, and further analyses were performed (Fig. 5a). Figure 5 and Figures S17 and S18 show the percent change from baseline in CCorr values for the two groups (post-needle insertion, post-needle manipulation, and post-needle withdrawal).

**Fig. 5.**
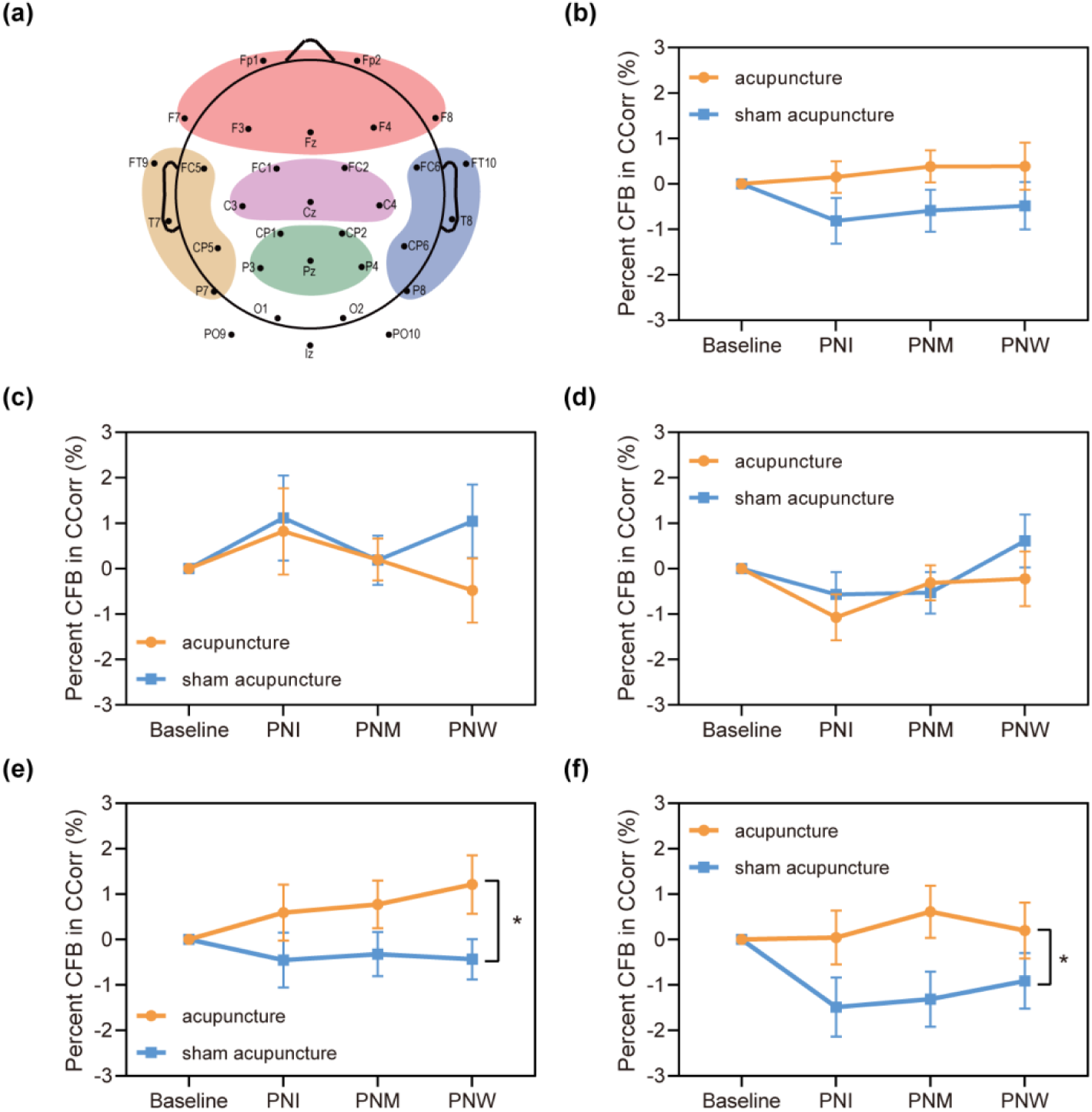
ROIs and change from baseline in CCorr in the fast-ripple frequency band in different regions. (a) The ROIs. The regions marked in the red area represent the frontal regions, the purple area represent the frontal-central regions, the green area represent the parietal regions, the brown area represent the left temporoparietal regions, and the blue area represent the right temporoparietal regions. (b) Percent CFB in CCorr in the frontal regions at baseline, PNI, PNM and PNW sessions. (c) Percent CFB in CCorr in the frontal-central regions at baseline, PNI, PNM and PNW sessions. (d) Percent CFB in CCorr in the parietal regions at baseline, PNI, PNM and PNW sessions. (e) Percent CFB in CCorr in the left temporoparietal regions at baseline, PNI, PNM and PNW sessions. (f) Percent CFB in CCorr in the right temporoparietal regions at baseline, PNI, PNM and PNW sessions. The stars indicate significant differences between the acupuncture and sham acupuncture groups. The data in (b)-(f) are plotted as the mean±SEM. Acupuncture, *n*=29; sham acupuncture, *n*=28. CFB, change from baseline; PNI, post-needle insertion; PNM, post-needle manipulation; PNW, post-needle withdrawal; SEM, standard error of the mean.

Acupuncture primarily affects the inter-brain connectivity in the fast-ripple frequency band in the left and right temporoparietal regions between the simulated patient and acupuncturist. ANOVA results revealed a significant main group effect in the left and right temporoparietal regions in the fast-ripple frequency band, and the percent change from baseline in the CCorr in the acupuncture group was significantly greater than that in the sham group (mixed-design ANOVA, left temporoparietal: F (1, 49) = 8.55, *p* = 0.005 and right temporoparietal: F (1, 55) = 4.49, *p* = 0.039) (Fig. 5e, f). Additionally, a significant main effect of the acupuncture session was found in the fast-ripple frequency band in the parietal regions (mixed-design ANOVA, F (3, 165) = 2.93, *p* = 0.035), and post-hoc tests revealed a significant increase in the post-needle insertion session compared with the baseline and post-needle withdrawal (two-tailed paired *t*-test, baseline vs. post-needle insertion: *p* = 0.024 and post-needle insertion vs. post-needle withdrawal: *p* = 0.016) (Fig. 5d). No significant differences or changes in the fast-ripple frequency band in the frontal and frontal-central regions (Fig. 5b, c) and the high-gamma and ripple frequency bands in the other ROIs were found (Figs. S17, 18).

## 4 DISCUSSION

This was the first hyperscanning study to investigate the inter-brain similarity and connectivity in HFOs across the brains of socially interacting individuals and the first EEG-based hyperscanning study to investigate the brain similarity, connectivity and synchronization between the simulated patient and acupuncturist during acupuncture.

Acupuncture practice pays attention to improve the psychological relationship between patients and acupuncturists, and rapport can be established even through physical contact alone during acupuncture^48^. The complete acupuncture process can be divided into needle insertion, needle manipulation and needle withdrawal. The patient and acupuncturist interact differently during these different sessions of acupuncture. The inter-brain similarity, connectivity and synchronization based on hyperscanning can identify brain mechanisms supporting the patient–acupuncturist relationship while this dyad is engaged in interactions. Using an EEG-based hyperscanning approach, this study performed patient-acupuncturist hyperscanning with simultaneous recordings of patient–acupuncturist interactions during different acupuncture sessions and examined the influence of different patient–acupuncturist interactions on the inter-brain similarity and connectivity between the patient and acupuncturist under acupuncture and sham acupuncture conditions.

In the present study, we developed an EEG hyperscanning method for simultaneous and objective recording of patient–acupuncturist interaction activities. In many studies, brain activity is commonly acquired together with data on behaviour^49–51^. The behaviour of patients and acupuncturists during acupuncture is difficult to record objectively given the complexity of the acupuncture process. In addition, previous platforms for acupuncture research based on hyperscanning have not focused on the simultaneous and objective recording of the behaviour of patients and acupuncturists during acupuncture^11,13,15^. Therefore, we used the E-prime program to objectify the behaviour of the acupuncturist (e.g., needle insertion and needle manipulation) as well as the behaviour of the patient (e.g., perception of needle insertion) and tagged them synchronously to the EEG data. Our method provides a platform for investigating the neural basis of patient–acupuncturist interaction-related EEG signals in a typical clinical setting.

The results revealed changes in inter-brain similarity and connectivity in the beta frequency band between the simulated patient and acupuncturist after needle insertion, and the changes in connectivity were observed mainly in the right temporoparietal regions. Moreover, the inter-brain connectivity of the fast-ripple frequency band in the left and right temporoparietal regions was significantly greater in the acupuncture group than in the sham acupuncture group. Finally, the inter-brain similarity of both the low-frequency bands and HFOs significantly decreased after both real and sham needle manipulation. These results are discussed below.

First, the neural mechanism involved in patient–acupuncturist interactions has been unclear. How the brains of the acupuncturist and patient work during patient‒ acupuncturist interactions in acupuncture remains unknown. A previous study suggested that INS in the PFC of the patient–acupuncturist dyad plays a critical role during needle manipulation^15^. A new finding in the current study was that the inter-brain similarity between the simulated patient and acupuncturist decreased at the global level (i.e., spatial correlation), but the inter-brain connectivity between them increased in the beta frequency band in the right temporoparietal region at the channel level (i.e., CCorr) after needle insertion. In addition, the results from further comparisons suggested that when the needle tip was subcutaneous, CCorr increased significantly in the acupuncture group. In addition, the increase was greater after needle insertion, whereas there was no change in the sham acupuncture group. The present observations provide a new perspective that neural synchronization in low-frequency bands is a potential neural mechanism for patient–acupuncturist interactions during acupuncture.

With respect to inter-brain connectivity at the channel level, enhanced connectivity of the beta frequency band has been found in many social settings, such as competition^52^ and face-to-face training^53^, and inter-brain synchrony is thought to be driven primarily by beta oscillations^49^. It has been proposed that beta frequency band synchrony might be related to the shared engagement and expectations of the peers’ actions^54,55^. In the present study, participants had expectations about acupuncture and the other participant’s actions. We found increased inter-brain connectivity in the beta band at the right temporoparietal region after needle insertion, potentially indicating that this activity is relevant for predicting each other’s actions and reactions after needle insertion and for predicting the effects of needle insertion. In addition, as shown by previous studies^56,57^, the increase in beta inter-brain connectivity after needle insertion may be related to mechanisms of awareness, intentionality and action planning experienced after needle insertion.

Moreover, in some studies of interpersonal interactions, such as face-to-face interactions^58^ and joint actions^59^, inter-brain synchronization is reflected mostly in the right temporoparietal region. This finding was confirmed in the present study. Research has shown that the right temporoparietal junction is a central node of the social brain^60^, which may account for the change in patient‒acupuncturist inter-brain connectivity in the right temporoparietal region in this study. However, EEG signals were only obtained from scalps rather than from cortices; therefore, brain activity in the right temporoparietal junction was not directly obtained. The next step could be to collect other brain data from participants for further source analysis.

The change in inter-brain connectivity at the global level contrasted with that at the channel level in the present study. This finding suggests that mechanisms of changes in inter-brain connectivity after needle insertion may differ at the global level and the channel level. The increased inter-brain connectivity in the right temporoparietal region after needle insertion may lead to a decrease in connectivity in other regions, which might be reflected in an overall decrease at the global level.

Second, HFOs are mostly seen and play a role in the diagnosis and treatment of epilepsy. However, an animal study revealed that HFOs are involved in social interactions^61^; therefore, we hypothesise that acupuncture may affect HFOs. Although previous hyperscanning studies have focused mostly on the interaction mechanisms at low-frequency bands, our current study adds the change in HFOs as one of the research focuses. A novel observation of the present study is that acupuncture enhances inter-brain connectivity in the fast-ripple frequency band. However, no reason could explain this observation because no research has explored HFOs during human social interactions. Further investigations are needed to verify the role of the fast-ripple frequency band during acupuncture. Furthermore, in many studies, inter-brain synchronization of temporoparietal regions increased in social interactions^62,63^. It has been proposed that this activity is potentially related to positive affect, which is a type of nonverbal social behaviour^64^. In the fast-ripple frequency band, enhanced inter-brain connectivity occurred in the left and right temporoparietal regions during acupuncture, which is potentially associated with the positive effects of the simulated patient and acupuncturist during acupuncture. Unlike the results of inter-brain connectivity in the beta frequency band above, enhanced inter-brain connectivity in the fast-ripple frequency band occurred during all the acupuncturist‒patient interactions associated with acupuncture. In the present study, acupuncture produced more positive effects than did sham acupuncture. Overall, we hypothesise that the fast-ripple frequency band may be more sensitive to positive affect and feedback between the patient and acupuncturist than the beta frequency band is.

Third, we applied spatial correlation, which is often used to explain the correlation between microstate prototype maps and their assigned EEG samples^65^, to observe changes in inter-brain similarity between the simulated patient and acupuncturist at the global level. A new finding of the current study was that the spatial correlations of both the low-frequency bands and HFOs significantly decreased after both real and sham needle manipulation. In the present study, a placebo effect was noted for both the true and sham needle manipulations. This observation suggests that the spatial correlation may primarily reflect the placebo effect rather than the specific effects of acupuncture. In both the acupuncture and sham acupuncture groups, needle manipulation was the most interactive session between the patient and acupuncturist, and this is potentially related to changes in spatial correlations. We hypothesised that spatial correlations were reduced after needle manipulation, possibly because the video training prior to the acupuncture increased the dyads’ expectations of *deqi* sensations. In addition, the interaction was the strongest during needle manipulation, which most closely matched the dyads’ expectations of *deqi* sensations; then, the expectations were reduced after needle manipulation.

A few study limitations should be mentioned. First, although this work provided the first analysis of three patient‒acupuncturist interactions during acupuncture, we did not allow for changes at multiple sessions during acupuncture, which should be further explored. Second, we only studied the EEG data using different methods and could not obtain other information about patient‒acupuncturist interactions. Thus, data such as electrocardiogram (ECG) data should be added to the analysis to explore the mechanisms and characteristics of patient‒acupuncturist interactions during acupuncture from multiple perspectives. In addition, stronger inter-brain synchronization in tasks may have required participants to have had some previous interactions, whereas the subjects in the present study had not been trained in the interaction task prior to the experiment. This might have affected the inter-brain similarity and connectivity. Moreover, only one acupuncturist was recruited in this study to ensure standardized acupuncture manipulation, and the experience of the acupuncturists might affect patient‒acupuncturist interactions. Finally, we only collected EEG from scalps rather than cortices. the EEG from scalps may not have originated from the cortex of the corresponding region; thus, other data, such as magnetic resonance imaging (MRI) data, could be collected for further source analysis.

## AUTHOR CONTRIBUTIONS

L.H., Y.G. and G.W. designed the study and participated in the data collection and analysis. S.J., Q.L., N.T., Y.Q., SJ.W., Q.Lyu., Y.L., Z.H., X.S., SY.W. and W.Z. did the trial and collected the data. S.J., Q.L. and N.T. contributed to data analyses. S.J. and Q.L. wrote the manuscript. Y.Q. participated in the manuscript revision. All authors contributed to the final manuscript.

## ACKNOWLEDGMENTS

We would like to thank Springer Nature Author Services for English language editing and Guillaume Dumas (main contributor of HyPyP) for the help in inter-brain connectivity analysis.

## DATA AVAILABILITY STATEMENT

The datasets generated and analysed during the current study are available in the the Open Science Framework repository, https://osf.io/z8cbw/?view_only=35c0ecd9e12948c9b2d626abbc8a4c7c.

## ETHICS APPROVAL AND CONSENT TO PARTICIPATE

This study was authorized by the Ethics Committee of the Institute of Acupuncture and Moxibustion, China Academy of Chinese Medical Sciences, with the approval number S2023-10-23-1.

